# Tailored Cell Cycle Modulation Enhances AAV Manufacturing: Balancing Arrest with Adaptive Stress Responses

**DOI:** 10.64898/2026.01.19.700387

**Authors:** Junneng Wen, Justin Sargunas, Dylan Carman, Noam Greenshtein, Michael J. Betenbaugh

## Abstract

Recombinant adeno-associated virus (rAAV) vectors show therapeutic potential, but their biomanufacturing is limited by low yields and high costs. Host cell-cycle modulation is emerging as a promising strategy to enhance rAAV production. Two G₂/M phase-arresting small molecules, ABT-751, a microtubule inhibitor, and helenalin, a thiol-reactive sesquiterpene lactone, were applied post-transfection in HEK293 cells to evaluate how cell-cycle arrest and stress pathways influence rAAV yields. ABT-751 induced G₂/M arrest with minimal cytotoxicity, leading to a near five-fold increase in rAAV vector genomes across multiple serotypes and production platforms. Helenalin caused G₂/M arrest, yet suppressed rAAV production. Comparative transcriptomic profiling (RNA-Seq) revealed that helenalin altered expression of a widespread set of genes (4,579) compared to control, characterized by rampant p53, ferroptosis, and endoplasmic reticulum dysregulation that overflowed into unfolded protein response with CHOP induction and apoptosis. ABT-751 elicited a more moderate, targeted response (1,895 differentially expressed genes) in a similar subset of pathways, including compensatory mechanisms mitigating oxidative stress. Together, these findings indicate that cell-cycle arrest alone is insufficient to improve rAAV yield. Indeed, tailored cell-cycle modulation, coupled with balanced activation of cellular stress pathways, can enhance rAAV manufacturing efficiency, facilitating more scalable and cost-effective gene therapy production strategies for the future.

## 1 Introduction

Recombinant adeno-associated viruses (rAAVs) are small, replication-defective, nonpathogenic parvoviruses that package a single-stranded DNA genome of approximately 4.7 kb within an icosahedral capsid, making them among the safest and most versatile vehicles for in vivo gene delivery [1], [2], [3], [4]. There are over 700 distinct clinical programs in development; however, despite remarkable clinical successes, the biomanufacturing process of rAAV remains a significant challenge, contributing to high costs, limited accessibility, and constrained production capacity [5], [6]. Plasmid triple transfection remains the standard production method despite inherent variability [7], and scalability of this system in suspension culture continues to challenge the field [8]. Further, the field has suffered multiple setbacks, including halted clinical programs and facility closures, due to persistent manufacturing limitations [9], [10]. In the current landscape, approved rAAV therapies including Zolgensma (onasemnogene abeparvovec) and Hemgenix (etranacogene dezaparvovec) are priced in the range of millions of dollars [11], [12].

Chemical modulation of the host-cell cycle has emerged as a promising lever to boost rAAV yields, which can in turn alleviate high production costs. In HEK293 suspension cultures, the microtubule inhibitor nocodazole has been shown to induce a reversible G₂/M arrest and increase rAAV genomic titers by 2-to 3-fold when applied shortly after transfection [13], [14]. This mirrors observations in other viral systems where G_2_/M arrest enhances replication by suppressing antiviral responses [15]. In our own evaluation of cell-cycle inhibitors, we identified ABT-751, an orally bioavailable sulfonamide antimitotic that binds the colchicine site on β-tubulin and likewise has been shown in trials to arrest cells in G₂/M [16], [17]. Other small molecule G₂/M arrestors work through tubulin-independent mechanisms; helenalin, a sesquiterpene lactone known to trigger G₂/M arrest and apoptosis in activated T cells [18], is one such example. These alternate paths to create cell-cycle blockades have thus far been unexplored in relation to viral vector productivity. As we show in this work, arresting the cell-cycle is not the sole determinant of productivity.

In this study, we comprehensively investigate the mechanisms underlying cell cycle–mediated modulation of rAAV productivity, highlighting the essential role of cellular stress and homeostasis pathways beyond simple cell-cycle arrest. Our findings offer important insights for future optimization of viral vector manufacturing, potentially guiding the strategic use of cell-cycle modulators and related compounds to enhance rAAV yield and improve the overall scalability and cost-effectiveness of gene therapy production.

## 2 Materials and Methods

### 2.1 Cell culture

An AAV2 helper free packaging system (Cell Biolabs, Catalog number VPK-402) consisting of three plasmids, namely pAAV-RC2 vector (native promoters: p5 for Rep78/Rep68, p19 for Rep52/Rep40, and p40 for Cap proteins), pHelper vector, and pAAV-GFP control vector (CMV promoter), was used as the plasmid system for rAAV production in this study. When different serotypes were tested, pAAV-RC2 was substituted. For generation of AAV5, pAAV2/5 was a gift from Melina Fan & Addgene Research Program (Addgene plasmid # 232922; http://n2t.net/addgene:232922; RRID:Addgene_232922). For generation of AAV9, pAAV2/9n was a gift from James M. Wilson (Addgene plasmid # 112865; http://n2t.net/addgene:112865; RRID:Addgene_112865). NEB 5-alpha competent E. coli cells were transformed using the obtained plasmids. The bacteria were amplified in LB media containing 100 μg/mL of ampicillin (Sigma Aldrich, Catalog number A9518) and glycerol stocks of selected colonies were generated by diluting the overnight cultures in a 1:1 ratio with 50% glycerol (Millipore Sigma, Catalog number G5516) and stored at −80°C. The selected colonies were also subjected to a diagnostic screen using a restriction enzyme digestion followed by an DNA agarose gel electrophoresis to verify for the correct plasmid size. The clones that passed the diagnostic test were then used for bacterial inoculation for the study. The plasmids were extracted and purified using QIAprep spin miniprep kit (QIAGEN, Catalog number 27104). For cell culture studies, plasmids were generated using the PureLink™ HiPure Expi Plasmid Gigaprep (Invitrogen, Catalog number K210009XP). The quality and concentration of the plasmids was measured using A260/280 using Nanodrop (Thermo Scientific, Catalog number ND-ONE-W).

A suspension adapted HEK293 host cell line, Expi293F (Gibco, Catalog number A14527), was used for this study. The cell growth medium used was the FreeStyle F17 expression medium (Gibco, Catalog number A1383501) and was supplemented with 4 mM GlutaMAX supplement (Gibco, Catalog number 35050061). Cells were thawed into 30 mL of prewarmed FreeStyle F17 medium containing 4 mM GlutaMAX in 125 mL plain-bottom shake flasks (Diagnocine, Catalog number NST-781101) in a humidified Multitron incubator (Infors HT, Product number SM100116-RH), at 37°C and 8% CO_2_ at 125 rpm. Cells were sub-cultured every 3-4 days when they reached a cell density of >3 × 10^6^ cells/ mL.

### 2.2 rAAV production via transient transfection

For a 30 mL rAAV production via co-transfection of the three AAV plasmids (triple transfection), the Expi293F cells were inoculated in prewarmed FreeStyle F17 medium supplemented with 4 mM GlutaMAX at a seeding cell density of 1.11 × 10^6^ cells/mL in 125 mL plain-bottomed shake flasks with a working culture volume of 27 mL on the day of transfection. This was done to obtain a final viable cell density of 1 × 10^6^ cells per mL after addition of the transfection cocktail.

The transfection cocktail was then prepared in a 15 mL sterile conical tube (Sarstedt, Catalog number 62.554.502). The three AAV plasmids were carefully added to the tube at a fixed molar ratio of 1:2:1 (pAAV-GFP: pAAV-RC2: pHelper). A total of 30 μg of plasmid DNA (pDNA) was added in order to target a final concentration of 1 μg/mL upon addition to the transfection flask. This translates to a ratio 1 μg of total pDNA per million cells in the culture. 3 mL of prewarmed fresh medium (FreeStyle F17 with 4 mM GlutaMAX) was then added to the conical tube containing the plasmids. A premade 1 mg/mL stock solution of linear polyethyleneimine hydrochloride (MW 40,000), also known as PEI Max (Polysciences, Catalog number 24765), was added to the cocktail at a PEI: pDNA w/w ratio of 2.2:1. In other words, 66 μL of the stock solution was added to the tube that already contained 30 μg of pDNA. The contents of the tube were thoroughly mixed by inverting the tube 3-5 times and incubated at room temperature for 7 minutes. After 7 minutes, the transfection cocktail was pipetted dropwise with continuous shaking into the transfection flask containing cell suspension.

The transfection flask was returned to a humidified Forma™ Direct Heat CO2 Incubator (Thermo Scientific, Catalog Number 360), at 37°C and 8% CO2 at 120 rpm. This study was done in biological triplicate or quadruplicate, and daily cell culture samples were collected for monitoring cell growth, transcriptomic profiling, and other attributes. On day 3 of the culture, i.e., 72 hours post-transfection, 1 mL aliquots of cell culture samples were frozen at −80°C for rAAV genome titer measurement at a later time.

### 2.3 Cell growth and viability

Cell culture samples were collected after every 24 hours for cell density and viability measurement during the cell culture process. Cells were stained and diluted using a 0.2% trypan blue solution (Gibco). Viable cell density (VCD) was used to determine the cell growth and was measured using Countess 3 (Invitrogen, Catalog number A49862). The cell culture viability was tracked using the trypan blue dye exclusion method.

### 2.4 AAV Genome Quantification by Digital PCR

Harvested AAV-containing culture was first subjected to sonication using a Branson SFX150 Sonifier with Handheld Converter (Branson Ultrasonics, Catalog number SFX150) at 20% power with 1-second on/off pulses for 2 minutes) to disrupt viral capsids and release genomes. Sonicated samples were then processed using the CGT Viral Vector Lysis Kit (QIAGEN, Catalog number 250272) according to the manufacturer’s instructions, which included DNase I digestion to remove non-encapsidated DNA, followed by chemical lysis steps and dilution in CGT Dilution Buffer to achieve the target copy concentration for dPCR.

Absolute quantification of AAV genomes was carried out on a QIAcuity 8.5K digital PCR system (QIAGEN, Catalog number 911012) using QIAcuity Nanoplate 8.5K 24-well (QIAGEN, Catalog number 250195) and the QIAcuity EG PCR Kit (QIAGEN, Catalog number 257520). Reactions were assembled and run strictly per the manufacturer’s protocol, and data were analyzed using the QIAcuity Software Suite with default Poisson-correction settings.

### 2.5 Cell Cycle Analysis by Flow Cytometry

Cells were harvested and counted to obtain approximately 3 × 10⁶ cells per condition. Cells were then washed twice with phosphate-buffered saline, or PBS (Gibco, Catalog number 10010023) at 300 × g and subsequently resuspended to achieve a final concentration of 3 × 10⁶ cells/mL. One milliliter of cell suspension was transferred into a 15 mL polypropylene, V-bottom tube (Sarstedt, Catalog number 62.554.502). Cells were fixed by the dropwise addition of 5 mL ice-cold 70% ethanol while gently vortexing to prevent cell clumping. Fixed cells were incubated for at least 1 hour at 4°C, after which they were either stored at −20°C for up to two weeks or immediately processed further.

Post-fixation, cells were pelleted by centrifugation at 800 × g for 5 minutes, followed by two washes in PBS. Pelleted cells were resuspended in 0.5 mL of FxCycle™ PI/RNase Staining Solution (Invitrogen, Catalog number F10797), mixed thoroughly, and incubated for 15-30 minutes at room temperature protected from light. Flow cytometric analysis was performed using a FACSCanto flow cytometry system (BD Biosciences). Cell cycle data were analyzed using FlowJo software (BD Biosciences) and fitted to the Dean-Jett-Fox model when possible. In cases where cell cycle inhibition precluded accurate model fitting, phases of the cell cycle (G_1_, S, G_2_/M) were manually determined by visual inspection and gating based on DNA content histograms.

### 2.6 Specific Productivity Analysis

To evaluate viral production efficiency independent of cell growth kinetics, Specific Productivity (Q_p_) was calculated as the total vector genomes (vg) divided by the total viable cells (TVC) at the time of harvest (72 h). This metric normalizes the viral yield against the viable biomass, isolating the cellular manufacturing performance from variations in cell proliferation rates caused by the cytostatic treatments.

### 2.7 RNA-Seq Library Preparation, Sequencing, and Primary Data Processing

Total RNA was extracted from treated HEK293 cells (ABT-751, helenalin, and DMSO control) using the Quick-RNA Miniprep Kit (Zymo Research, Catalog number 11-330), following the manufacturer’s protocol. Samples were shipped on dry ice to Novogene (Sacramento, CA), where RNA integrity and concentration were assessed using an Agilent 2100 Bioanalyzer (RIN ≥8) and a Qubit RNA HS Assay, respectively. At Novogene, poly(A) RNA was captured with oligo(dT) magnetic beads and fragmented by divalent cations under elevated temperature. First-strand cDNA synthesis was performed using random hexamers, followed by second-strand synthesis. End repair, A-tailing, and ligation of Illumina P5/P7 adapters were carried out, and fragments of 200–350 bp were size-selected. Libraries were quantified by Qubit dsDNA HS Assay and qPCR (KAPA Library Quantification Kit), and size distributions were confirmed on the Bioanalyzer. Cluster generation was performed on a NovaSeq X Plus S4 flow cell, and sequencing was run as 150 bp paired-end using NovaSeq X Plus chemistry on an Illumina NovaSeq X Plus platform, generating 121.1 Gb of raw data (mean ∼30 million read pairs per sample). Base calling and demultiplexing were performed with Illumina bcl2fastq software. Raw reads were filtered to remove adapter sequences, reads containing >10% ambiguous bases (N), or reads with >50% of bases having Phred score ≤5. Clean reads were retained at 96.9–97.9% per sample. Overall error rates were ≤0.02%, Q20 scores ≥99.0%, Q30 scores ≥96.1%, and GC content ranged from 51.8–53.2% Resulting clean FASTQ files were used for downstream alignment and expression quantification.

### 2.8 Sequencing Data Processing and Gene-Level Quantification

Adapter trimming and quality assessment were performed on the clean FASTQ files using Cutadapt (v3.4) [19] to remove standard Illumina TruSeq P5/P7 adapter sequences (3′-AGATCGGAAGAGCACACGTCTGAACTCCAGTCAC; 5′-GTTCAGAGTTCTACAGTCCGACGATC), followed by FastQC (v0.11.9) [20] to confirm adapter removal and overall read quality. Trimmed reads were aligned to the human reference genome GRCh38 with HISAT2 (v2.2.1) [21] using the--rna-strandness RF option to preserve strand specificity. Resulting SAM files were converted to BAM, sorted, and indexed using SAMtools (v1.12) [22]. Finally, gene-level quantification was carried out with featureCounts (Subread v2.0.1) [23] against GENCODE v38 annotations, employing the -p flag for paired-end data and -s 2 for reverse-strand libraries to generate the raw gene count matrix for downstream differential expression analysis.

### 2.G Differential Expression Analysis

Raw count matrices were imported into R (v4.2.2) and analyzed with DESeq2 (v1.36.0) [24]. Low-count genes (total count ≤0 across all samples) were removed prior to normalization. The design formula ∼ condition contrasted each treatment (ABT-751, Helenalin) against DMSO, and an additional head-to-head contrast of ABT-751 vs Helenalin was performed. Size factors and dispersion estimates were calculated with default methods, and log₂ fold-changes were shrunken using the apeglm method (apeglm v1.18.0) [25] to improve effect-size estimates for low-count genes. Genes with Benjamini–Hochberg adjusted p-value (padj) < 0.05 were considered significant.

### 2.10 Gene Ontology (GO) and Pathway Enrichment

Significantly up- or down-regulated gene lists (padj < 0.05) were mapped from HGNC symbols to Entrez IDs via org.Hs.eg.db (v3.15.0) using AnnotationDbi’s mapIds(). Genes without valid Entrez ID mappings were excluded from downstream pathway enrichment analyses. GO enrichment (Biological Process ontology) was performed with clusterProfiler (v4.6.0) [26], [27] using a Benjamini– Hochberg correction and an adjusted p-value cutoff of 0.05. Enriched GO terms were visualized as dot plots with enrichplot (v1.16.0).

KEGG pathway enrichment was similarly conducted via clusterProfiler’s enrichKEGG() function against Homo sapiens annotations, filtering terms with padj < 0.05. Disease- and neural-related pathways were excluded by keyword (e.g. “cancer”, “neuro”) to focus on cellular processes. The top 10 enriched KEGG terms per comparison were plotted using dotplot().

## 3 Results

### 3.1 G₂/M Arrest by ABT-751 Dramatically Boosts AAV2 Production

To assess the impact of cell-cycle modulation on recombinant AAV production, we transiently triple-transfected HEK293 cells with packaging, transgene, and helper plasmids in the presence or absence of a small-molecule cell-cycle modulator, ABT-751 (**Fig. 1**). We first monitored viable cell density (VCD) and viability in Expi293F HEK293 cultures transfected for AAV2 and treated with either ABT-751 or vehicle (DMSO) (**Fig. 2A**). Vehicle-treated cultures expanded rapidly, reaching ∼4.0 × 10^6^ cells/mL by 72 h post-transfection, whereas ABT-751 limited proliferation, with VCD plateauing at ∼1.2 × 10^6^ cells/mL. While the viability of the control group stayed above 95% throughout, ABT-751– treated cultures showed a modest decline beginning at ∼24 h but remained above ∼65% viable over 3 days of culturing.

**Figure 1:**
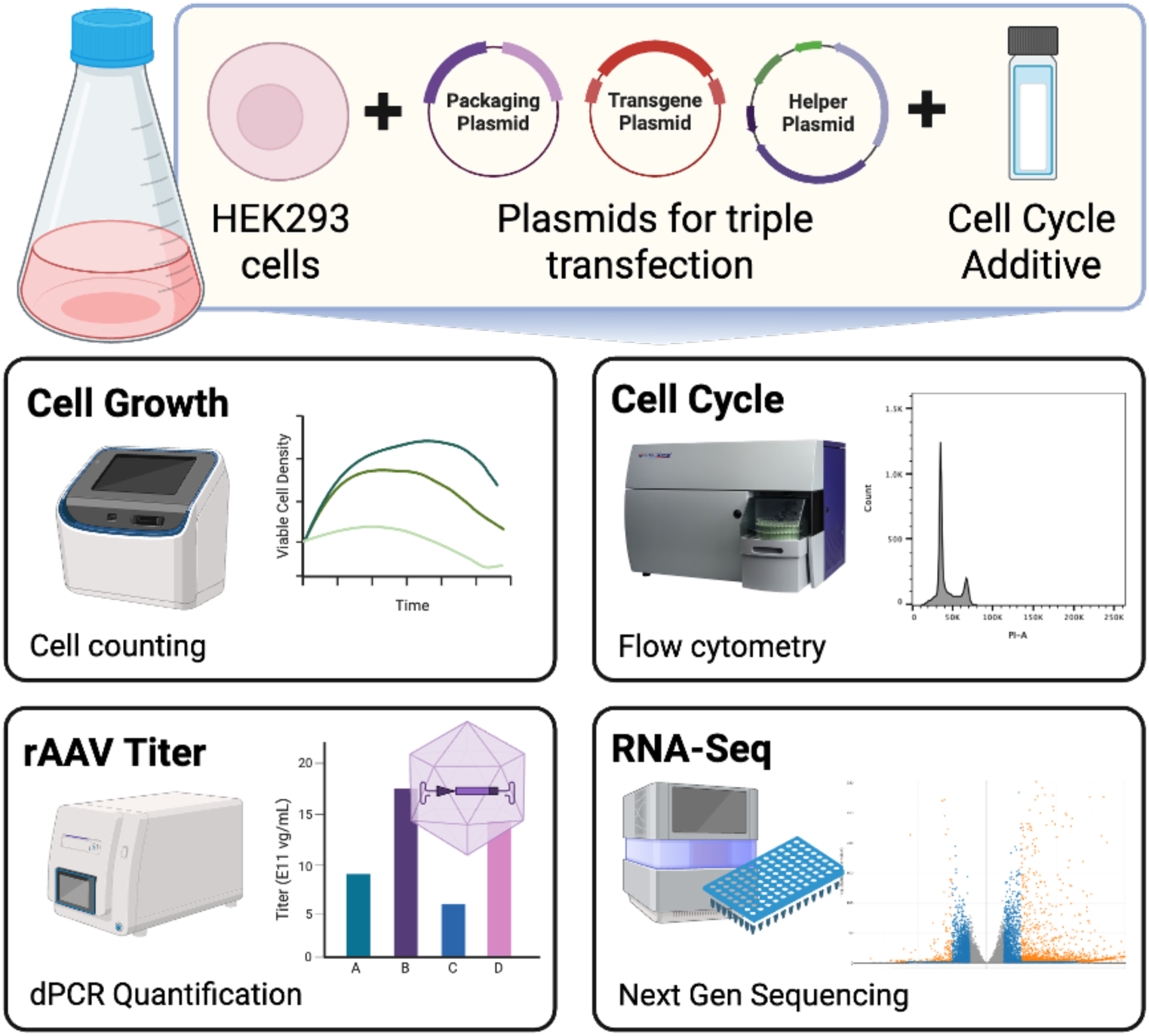
Schematic of the cell-cycle modulation workflow during rAAV production. HEK293 cells were transiently triple-transfected with packaging, transgene, and helper plasmids in the presence or absence of a small-molecule cell-cycle additive. Post-transfection, (i) cell growth and viability were tracked via automated cell counting, (ii) cell-cycle phase distribution was assessed by flow cytometry, (iii) rAAV vector genomes were quantified by digital PCR to determine viral titer, and (iv) host-cell gene expression levels were profiled by next-generation sequencing. Created with BioRender.

**Figure 2:**
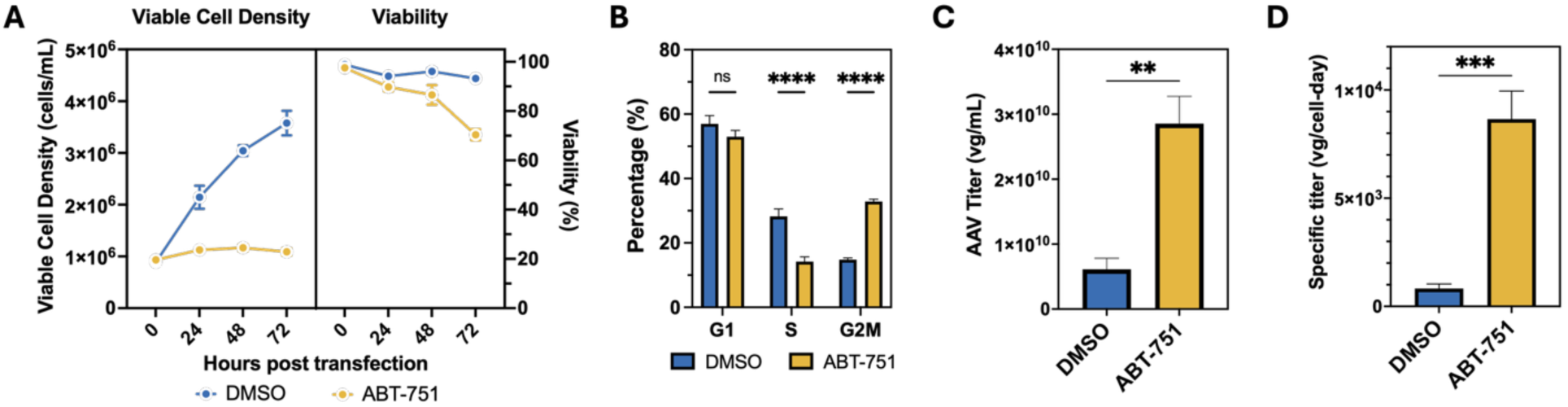
ABT-751 treatment enhances AAV2 production in HEK2G3 cells. **(A)** Viable cell density and viability of HEK293 cells treated with ABT-751 or vehicle (DMSO) post-transfection, shown as mean ± SD. **(B)** Quantification of cell cycle distribution showing the percentage of cells in G1, S, and G_2_/M phases. Statistical analysis was by two-way ANOVA with Sidák’s multiple-comparisons test: G₁, ns; S, **** p < 0.0001; G₂/M, **** p < 0.0001. **(C)** AAV2 titer (vector genomes per milliliter, vg/mL) measured by dPCR. Statistical significance was determined using an unpaired t-test (*p* = 0.001). **(D)** Specific AAV2 titer normalized to viable cell count and culture duration (vg/cell*day). Statistical significance was determined using an unpaired t-test (*p* = 0.0005). n = 3 biological replicates per condition. p-values: ** p < 0.01, *** p < 0.001.

As ABT-751 is a microtubule inhibitor, we first verified on-target engagement and quantified how much the cell-cycle distribution shifted by profiling DNA content via flow cytometry with PI staining (gating in **Supplemental Fig. S1**; distributions in **Fig. 2B**). Debris was excluded on forward/side scatter (FSC/SSC), doublets were removed by area–height/width discrimination, and PI-positive singlets were quantified for DNA content. The resulting histograms showed well-resolved G_1_ and G_2_/M peaks across replicates in both conditions, indicating that the observed phase shifts reflected biological differences rather than gating artifacts.

At 72 h post-transfection, compared to vehicle-treated cells, ABT-751 shifted the distribution from 57% to 53% G₁, reduced S from 28% to 14% (p = 9 × 10^-4^), and increased G₂/M from 15% to 33% (p = 4 × 10^-6^) (**Fig. 2B).** The remaining population, sub-G_1_, showed lower DNA content indicative of genome fragmentation and apoptosis (**data not shown**) [28]. Together, these results were consistent with microtubule-based mitotic blockade. We next measured total AAV2 vector genomes in culture bulk harvests by digital PCR (**Fig. 2C**). ABT-751–treated cells produced 2.8 × 10^10^ vector genomes (vg)/mL on average, nearly five-fold higher than vehicle controls (5.6 × 10^9^ vg/mL; p = 0.001; n = 3). Finally, when we normalized vector yield to viable cell count, ABT-751 treatment increased specific productivity from 7.5 × 10^2^ to 8.6 × 10^3^ vg/cell-day (p = 0.0005; **Fig. 2D**), approximately 11-fold higher. Together, these data demonstrate that ABT-751 slows HEK293 proliferation via G_2_/M arrest while substantially enhancing both total and specific AAV2 production.

### 3.2 ABT-751–Driven G₂/M Arrest Broadly Enhances AAV Production Across Serotypes and Cell Lines

To determine the optimal concentration of ABT-751 for boosting productivity while minimizing negative cell health effects, we conducted a dosage screening assay. We selected a maximum dose of 9.69 μM to align with the highest dose previously used on other mammalian cells lines [29], and tested concentrations down to 0.27 μM. Cells transfected for AAV2 production were treated immediately following transfection with four different ABT-751 doses, and cell stages for each condition were measured after three days (**Supplemental Fig. S2**). We observed an increase in the G_2_/M population from 17% to 35% between the doses of 0.27 μM and 2.69 μM ABT-751 at 72 h. Concentrations beyond 2.69 μM showed minimal impact on the cell cycle distribution. This preliminary screen established 2.69 μM ABT-751 as the maximum necessary dose for G_2_/M blockade.

To establish the minimal effective concentration for robust G₂/M arrest, HEK293 cells were treated with either 2.69 µM or 0.97 µM [29] ABT-751 immediately at transfection. Cell growth and viability were monitored daily; both treated samples demonstrated lower VCD, with the higher dose having a marginally harsher effect on cell viability by day 3 (**Supplemental Fig. S3A**). DNA content was also measured daily (**Supplemental Fig. S3B**). Both doses induced a clear, time-dependent accumulation in G₂/M and accompanying reduction in S-phase compared to the DMSO control. While the higher dose achieved near-maximal arrest by day 2 (G₂/M > 20%), the lower dose was sufficient to elicit a sustained G₂/M enrichment. Therefore, to preserve culture health and minimize chemical residuals in future downstream applications, we carried 0.97 µM forward for timing and productivity studies.

We next asked what dosing schedule of ABT-751 best balances cell growth and vector yield. Using the lower 0.97 µM dose, we added inhibitor at –6 h, 0 h, 24 h or 48 h relative to transfection (**Supplemental Fig. S3C–D**). All conditions with early addition of ABT-751 (−6 h or 0 h) showed reduced viability (65-70% on day 3) as compared to vehicle controls. However, AAV2 titers were highest when ABT-751 was present at the time of transfection and progressively decreased with later additions. These data indicate 0.97 µM ABT-751 added immediately following transfection (0 h) was the optimal regimen for inducing G₂/M arrest and maximizing AAV2 output.

To examine applicability across serotypes, we applied 0.97 µM ABT-751 to HEK293 cultures producing AAV2, AAV5, and AAV9 (**Fig. 3A–B**). For AAV2, bulk titer averaged 2.9×10¹⁰ vg/mL with ABT-751 compared to 6.1×10⁹ vg/mL with vehicle, and specific productivity averaged 2.6×10⁴ vg/cell-day compared to 1.7×10³ vg/cell-day. For AAV5, bulk titers were 1.3×10¹¹ versus 1.9×10¹⁰ vg/mL, and specific productivities were 1.4×10⁵ versus 8.2×10³ vg/cell-day. For AAV9, bulk titers were 1.5×10¹¹ versus 3.3×10¹⁰ vg/mL, and specific productivities were 1.5×10⁵ versus 1.6×10⁴ vg/cell-day. Across serotypes, these gains were significant by two-way ANOVA with Sidák correction (all p < 10⁻⁴).

**Figure 3:**
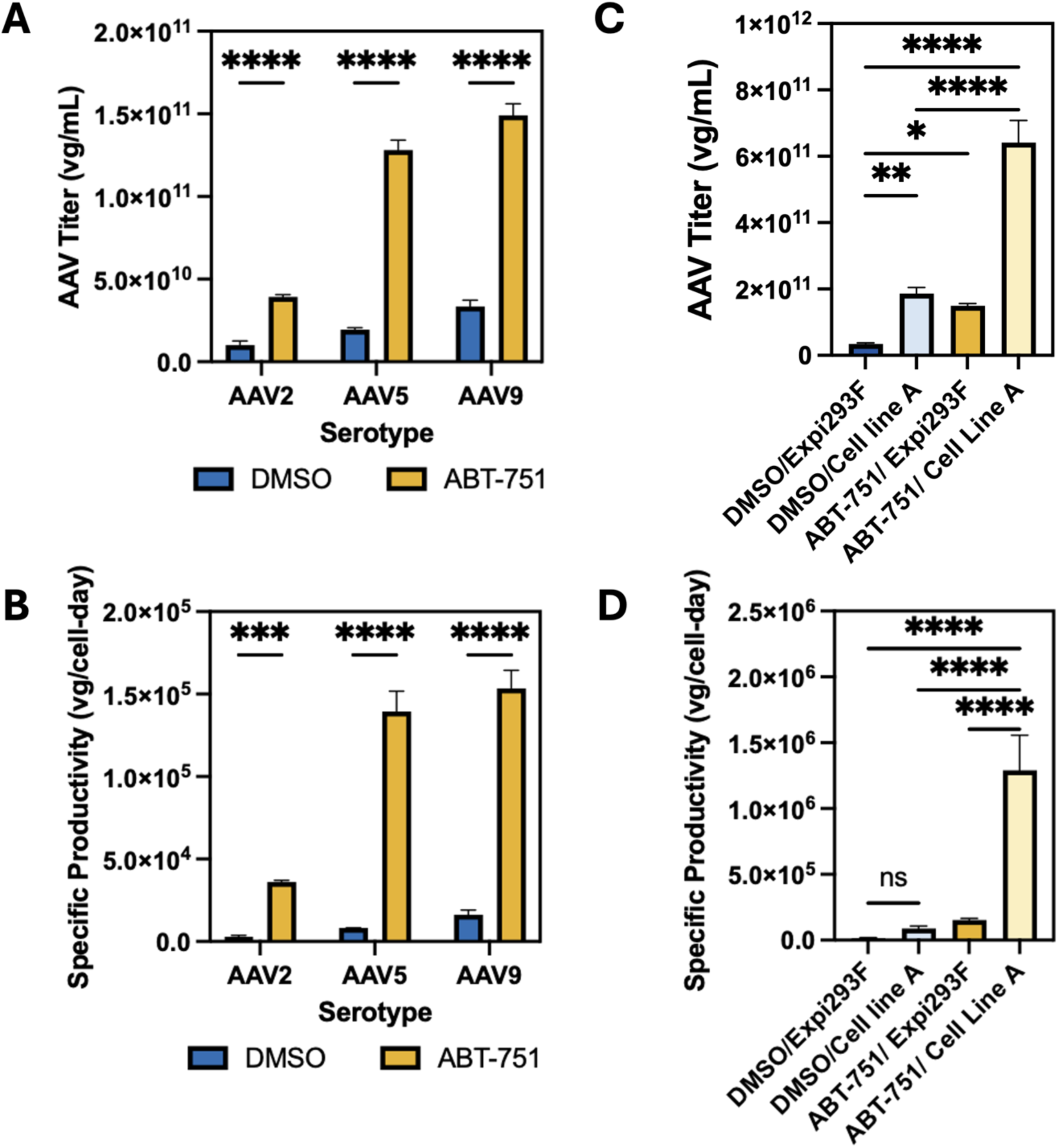
ABT-751–driven G₂/M arrest boosts AAV yields across serotypes and scales. **(A)** Bulk AAV titer (vg/mL) for three capsids (AAV2, AAV5, AAV9) produced in HEK293 cells treated with DMSO vehicle or 0.97 µM ABT-751. **(B)** Specific AAV productivity (vg·cell⁻¹) for the same serotypes and treatments. In (a–b), bars are mean ± SD (n = 3); two-way ANOVA (factors: serotype, treatment) with Sidák’s multiple comparisons: *p < 0.05; **p < 0.01; ****p < 0.0001; ns, not significant. **(C)** Bulk AAV titer in Expi293F and Cell Line A—± 0.97 µM ABT-751. **(D)** Specific productivity in the same four conditions. In (c–d), bars are mean ± SD (n = 3); one-way ANOVA with Tukey’s post-hoc test: *p < 0.05; **p < 0.01; ****p < 0.0001.

We then evaluated platform compatibility by comparing the Expi293F suspension cells with another suspension cell line derived from ATCC (Cell Line A; **Fig. 3C–D**). Both cell lines saw a significant increase in viral titer to near or above 1 ×10^11^ vg/mL in the presence of the additive with Expi293F and Cell Line A seeing 4.4-fold and 3.4-fold bulk titer increases, respectively (**Fig. 3C**). Similarly, specific AAV productivity increased 9.5-fold for Expi293F cells, and 14.5-fold for Cell Line A (**Fig. 3D**) with treatment effects being significant as measured by one-way ANOVA with Tukey adjustment (all p < 10⁻⁴). Collectively, ABT-751–induced G₂/M arrest delivered substantial gains in AAV production across multiple serotypes and different producer platforms, showing generalizability beyond one single platform.

### 3.3 Similar Growth and Cell-Cycle Profiles, Divergent rAAVG Yields under ABT-751 and Helenalin

To test whether the ABT-751 boost simply reflected the cell-cycle state in AAV9-producing HEK293 cultures, we screened a focused panel of cell cycle modulators: ABT-751, paeoniflorin [30], camptothecin (CPT) [31], and helenalin [18], alongside a DMSO vehicle and non-transfected controls. All compounds were tested at literature-supported concentrations. We quantified day 3 DNA content, culture health, and rAAV output (**Supplementary Fig. S4A-C**). Interestingly, ABT-751 and helenalin each produced pronounced G₂/M accumulation with S-phase depletion, and cultures reached similar viable cell density and viability at harvest. Despite comparable cell-cycle profiles, titers diverged: ABT-751 yielded high titers relative to DMSO, paeoniflorin offered modest positive improvement, whereas helenalin and CPT addition resulted in lower titers (**Supplementary Fig. S4D**). Thus, G₂/M arrest alone did not necessarily increase rAAV yield. The specific cellular programs engaged by each compound dictated their productivity, underscoring that not all cell-cycle inhibitors are functionally equivalent for rAAV manufacturing applications.

Motivated by the substantial titer divergence despite G₂/M enrichment, we focused on 0.97 μM ABT-751 and 5 μM helenalin for a head-to-head comparison to better characterize how different cell cycle inhibitors can affect HEK cell behavior and AAV production. We first compared viable cell densities across treatments at 0, 24, 48, and 72 h post-transfection (**Fig. 4A**). At 0 h and 24 h, there were no significant differences in VCD between DMSO, ABT-751, and helenalin (all p > 0.05). By 48 h and 72 h, both ABT-751- and helenalin-treated cultures showed significantly lower VCD than DMSO control. Viability followed a similar trend with reduced levels for both ABT-751 and helenalin relative to DMSO at 48 and 72 hours, indicating that the two compounds exerted similar impacts on general culture health.

**Figure 4:**
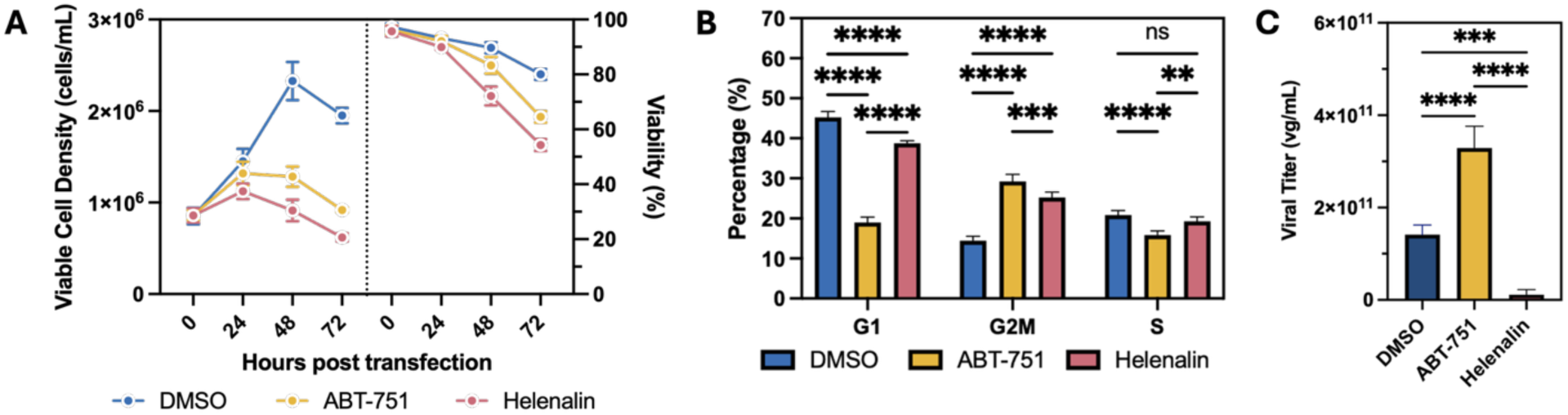
Differential effects of helenalin and ABT-751 on HEK2G3 rAAVG production. **(A)** Viable cell density (cells/mL, left axis) and cell viability (%; right axis) of HEK293 cells treated with 0.1% DMSO (blue circles), 0.97 µM ABT-751 (yellow circles) or 5 µM helenalin (pink circles), measured at 0, 24, 48 and 72 h post-transfection. Data are mean ± SD (n = 4). **(B)** Cell-cycle phase distribution of HEK293 cells harvested at 72 h post-transfection under the same treatment conditions. Bars represent the percentage of cells in G₁, G₂/M, and S phases for DMSO (blue bars), helenalin (pink bars), and ABT-751 (yellow bars); mean ± SD (n = 3). Statistical significance by two-way ANOVA with multiple comparisons: ****p < 0.0001, ***p < 0.001, **p < 0.01, ns = not significant. **(C)** rAAV9 vector genome titer in culture supernatants at 72 h post-transfection, determined by digital PCR. Bars show mean ± SD (n = 4) with individual replicates overlaid. Statistical significance by one-way ANOVA with Tukey’s post hoc test: DMSO vs ABT-751, ****p < 0.0001; DMSO vs helenalin, ***p < 0.001; ABT-751 vs helenalin, ****p < 0.0001.

To determine whether these agents impose distinct cell-cycle profiles, we quantified DNA content at 72 h (**Fig. 4B**). Both treatments significantly increased G₂/M populations relative to DMSO (14% G₂/M): ABT-751 to 29% G₂/M (p < 0.0001) and helenalin to 25% G₂/M (p < 0.0001). ABT-751 also significantly reduced S phase from 20% to 17% (p < 0.0001), while helenalin showed no significant effect on S phase. Compared to DMSO control (47% G₁), both treatments reduced the G₁ population: ABT-751 to 18% G₁ (p < 0.0001) and helenalin to 39% G₁ (p < 0.0001). As with previous cell cycle quantifications, the remaining cells were grouped into a sub-G_1_ population (**data not shown**). Although the G_1_ population was dramatically lower in the ABT-751 treatment versus helenalin with S and G_2_/M at similar levels, this does not necessarily indicate a significantly larger apoptotic population. Apoptosis tracking via DNA measurement is not a specific or quantitative measure of total apoptosis; the stage of apoptosis, extent of DNA fragmentation, chromatin state, cell morphology, and several other factors limit applicability of interpreting the sub-G_1_ peak [32], [33], [34]. As such, these data confirm that both compounds effectively arrest cells in G₂/M with similar impacts on cell health.

Interestingly, digital PCR quantification of rAAV9 genomes in lysed harvests at 72 h revealed that ABT-751 elicited a consistent ∼3-fold bulk titer increase over DMSO, whereas helenalin severely suppressed production relative to the DMSO control (**Fig. 4C**). Together, these results show that although both compounds perturb the cell cycle and slow growth in AAV9-producing HEK293 cultures, ABT-751 enhances vector yield while helenalin treatment is detrimental to AAV titers.

While ABT-751 treatment resulted in a lower final viable cell density (VCD) due to the induced cell cycle arrest, the volumetric titer was significantly increased. When normalized for cell number, the specific productivity (Q_p_) of ABT-751 treated cells was approximately 10-fold higher than DMSO controls. This indicates that the treatment transforms the host cells into highly efficient viral factories, prioritizing capsid production over biomass accumulation. Conversely, helenalin treatment resulted in a loss of specific productivity, correlating with the onset of cellular toxicity.

### 3.4 Global RNA-Seq analysis reveals divergent transcriptional programs induced by ABT-751 and helenalin

To dissect mechanisms beyond cell-cycle position, we performed RNA-seq at 72 h in AAV9-producing HEK293 cultures treated with ABT-751, helenalin, or DMSO (**Fig. 5**). Principal component analysis of variance-stabilized counts (**Fig. 5A**) revealed that helenalin-treated replicates separate sharply from both DMSO control and ABT-751 especially along PC1 (81% of variance), while PC2 (3%) captures the subtler shift induced by ABT-751 relative to DMSO control. GSEA analysis of PC1 gene loadings (**Supplementary Table S1**) revealed that this major axis of variation primarily reflects coordinated suppression of mitochondrial energy metabolism, including oxidative phosphorylation, aerobic respiration, and electron transport chain processes (all NES < −3.4, p < 1×10⁻⁶), along with activation of cellular stress responses, autophagy, and cell cycle regulation pathways in the helenalin treated samples. Thus, helenalin induces more severe metabolic disruption than ABT-751, explaining their separation along PC1.

**Figure 5:**
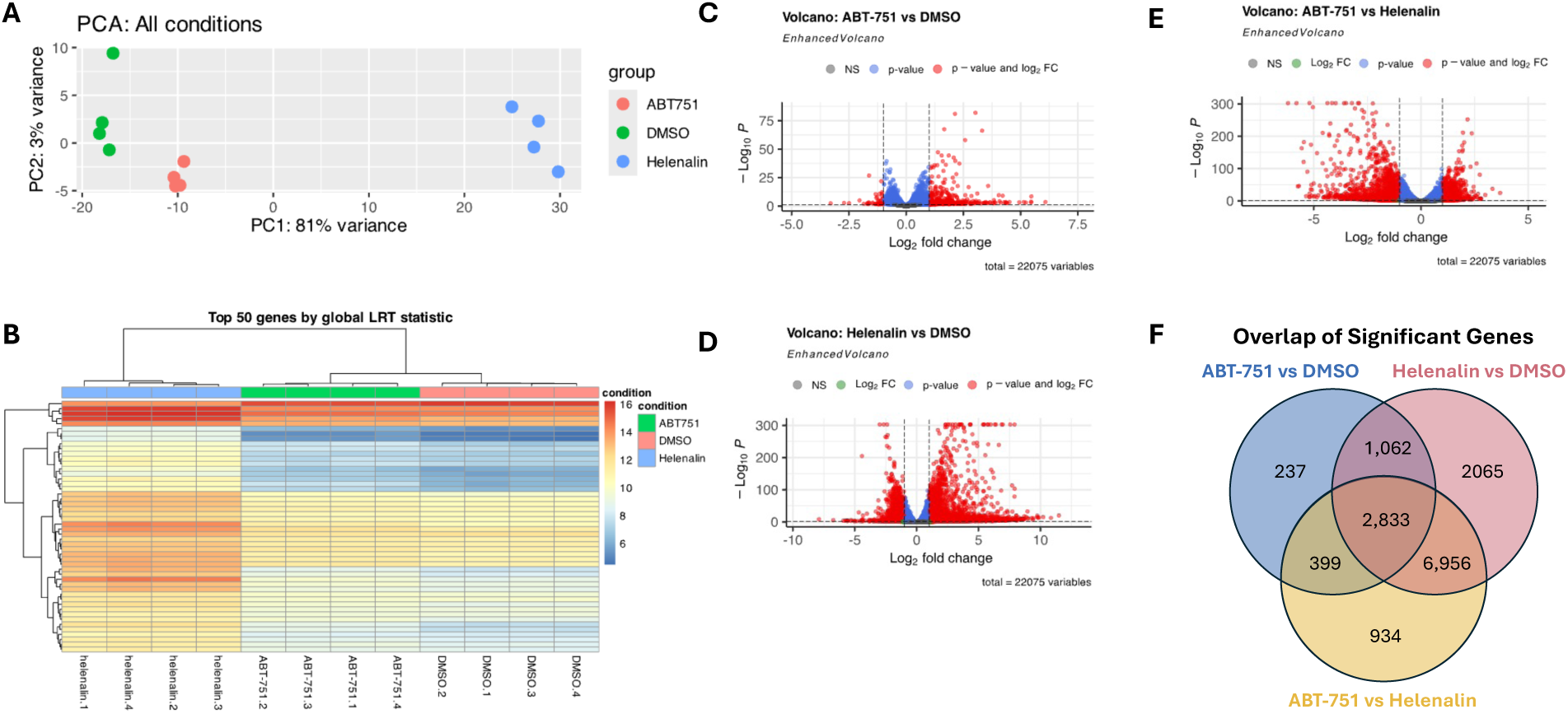
Global transcriptomic responses to ABT-751 and helenalin in HEK2G3 cells. **(A)** Principal component analysis of variance-stabilized counts for four replicates each of DMSO (green), ABT-751 (red) and helenalin (blue) treatments. PC1 (81% variance) separates helenalin from the other groups, while PC2 (3%) distinguishes ABT-751 from control. **(B)** Heatmap of the top 50 genes by likelihood-ratio test across all three conditions. Rows are genes (z-scored expression), columns are individual samples, and clustering was performed with Euclidean distance. **(C)** Volcano plot for ABT-751 vs DMSO: log₂ fold change (x-axis) versus –log₁₀(padj) (y-axis). Grey points are non-significant (padj ≥ 0.05), blue points meet padj < 0.05 only, and red points meet both padj < 0.05 and |log₂FC| ≥ 1. Total tested genes = 22,075; significant genes = 4,531. **(D)** Volcano plot for helenalin vs DMSO, plotted as in (c); significant genes = 12,916. **(E)** Volcano plot for ABT-751 vs helenalin, plotted as in (c); significant genes = 11,122. **(F)** Venn diagram of significant genes (padj < 0.05) in each contrast. Counts indicate genes unique to or shared among ABT-751 vs DMSO (4,531 total; 237 unique), helenalin vs DMSO (12,916 total; 2,065 unique), and ABT-751 vs helenalin (11,122 total; 934 unique), with 2,833 genes common to all three.

Hierarchical clustering of the 50 most variable genes by likelihood-ratio test (**Fig. 5B**) resolved three clusters: a helenalin-specific induction module, an ABT-751-repressed module, and a set with control-like expression. Complete differential-expression tables for each contrast are provided in **Supplementary Table S2**, including statistics for all genes (padj values and log₂ fold-changes).

Pairwise differential expression (padj < 0.05, |log₂FC| ≥ 1) identified 4,531 genes whose expression was significantly altered by ABT-751 versus DMSO (**Fig. 5C**) and a substantially larger set of 12,916 genes dysregulated by helenalin versus DMSO (**Fig. 5D**). These counts reflect all statistically significant DEGs prior to functional filtering and were used for global comparisons, clustering, and Venn diagram analyses. Direct comparison of ABT-751 to helenalin uncovered 11,122 genes with significant up- or down-regulation (**Fig. 5E**), confirming divergent modes of action. Venn diagram analysis of all significantly dysregulated genes (padj < 0.05; **Fig. 5F**) showed 2,833 genes common to every contrast (core stress response), 1,062 genes shared only between each compound and control, and discrete gene sets unique to ABT-751 (237 genes), helenalin (2,065 genes), or the ABT-751 versus helenalin head-to-head comparison (934 genes). These data demonstrate that, while both agents engage shared adaptive pathways, helenalin provokes a markedly broader transcriptional reprogramming than ABT-751.

To explore the biological processes and pathways underlying the transcriptional responses to ABT-751 and helenalin, we performed Gene Ontology (GO) Biological Process and KEGG pathway enrichment analyses. For pathway-level interpretation, the full DEG lists (padj < 0.05; **Supplementary Table S2**) were filtered to genes with valid Entrez ID mappings and sufficient annotation for functional enrichment, yielding 1,895 genes for ABT-751 and 4,579 genes for helenalin. These functionally annotated subsets were used for GO and KEGG enrichment analyses shown in **Fig. 6**.

**Figure 6:**
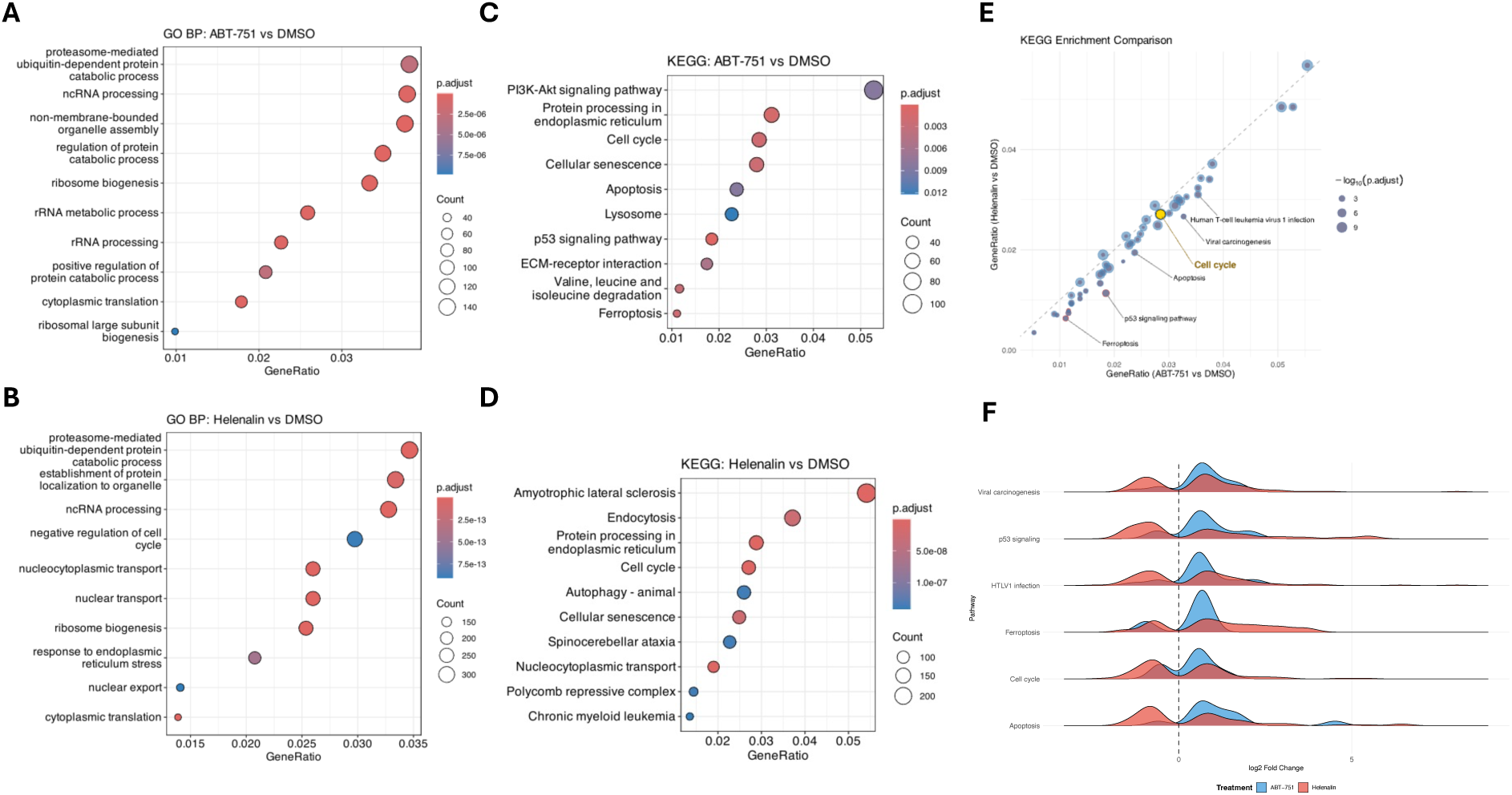
Pathway enrichment analysis reveals divergent stress response programs induced by ABT-751 and helenalin in AAVG-producing HEK2G3 cells. **(A)** Gene Ontology Biological Process (GO BP) enrichment for ABT-751 vs DMSO at 72 h post-transfection. Dot size indicates gene count; color intensity represents adjusted p-value significance. ABT-751 primarily enriches protein quality control and translational processes. **(B)** GO BP enrichment for helenalin vs DMSO. Helenalin induces broader transcriptional reprogramming, with enrichment of stress responses, nucleocytoplasmic transport, and cell cycle regulation pathways. **(C)** KEGG pathway enrichment for ABT-751 vs DMSO. Notable enrichment includes PI3K-Akt signaling, protein processing in endoplasmic reticulum, cell cycle, and ferroptosis pathways. **(D)** KEGG pathway enrichment for helenalin vs DMSO. Helenalin shows distinct enrichment of neurodegeneration-associated pathways (amyotrophic lateral sclerosis, spinocerebellar ataxia), autophagy, and chromatin regulation (polycomb repressive complex). **(E)** Cross-condition KEGG enrichment comparison plotting ABT-751 vs DMSO (x-axis) against helenalin vs DMSO (y-axis) gene ratios. Points below the diagonal indicate stronger enrichment under ABT-751. Highlighted pathways show the largest treatment divergences, with ABT-751 showing concentrated enrichment in checkpoint and stress-related pathways. **(F)** Expression distribution comparison across key pathways for ABT-751 (blue) and helenalin (red) treatments. Density plots show log₂ fold changes for genes within each pathway, demonstrating helenalin’s broader and more extreme transcriptional responses compared to ABT-751’s more focused effects.

We mapped differential expression onto the KEGG Cell cycle pathway using Pathview (**Supplementary Fig. S5**). Cell cycle pathway analysis revealed that both compounds affected the same regulatory nodes but with markedly different intensities. Helenalin triggered stronger activation of G_2_/M checkpoint kinases (*WEE1*, *CHEK1*) that prevent mitotic entry, and more complete suppression of the APC/C complex (*CDC20*, *CDC27*) required for mitotic exit. In contrast, ABT-751 showed more selective effects, retaining partial expression of S/G_2_ cyclins (*CCNA2*) and G_1_/S cyclins (*CCNE1*). These patterns (**Supplementary Table S3**) indicate that while both compounds arrest cells at G_2_/M, helenalin enforces multiple checkpoint barriers more stringently, consistent with its broader transcriptional disruption.

In the ABT-751 versus DMSO comparison (**Fig. 6A,C**), GO biological process enrichment centered on protein synthesis and quality control (proteasome-mediated catabolism, ribosome biogenesis, cytoplasmic translation) and RNA processing pathways. KEGG analysis revealed enrichment primarily in stress response (p53 signaling, apoptosis, cellular senescence) and protein processing pathways (ER protein processing, lysosome function), with additional metabolic adaptations (PI3K-Akt signaling, ferroptosis) (**Supplementary Table S4**).

Helenalin versus DMSO comparison (**Fig. 6B,D**) showed substantially broader pathway disruption (4,579 DEGs vs 1,895 for ABT-751). While GO terms overlapped with ABT-751 in proteolysis pathways, helenalin uniquely enriched nuclear transport dysfunction (nucleocytoplasmic transport, nuclear export) and severe ER stress responses. KEGG analysis revealed helenalin-specific enrichment of neurodegeneration pathways (amyotrophic lateral sclerosis, spinocerebellar ataxia) and cellular trafficking disruption (endocytosis, autophagy) absent in ABT-751 samples (**Supplementary Table S4**).

To identify the specific genes driving pathway enrichments, we performed detailed gene-level analysis of key enriched pathways, including p53 signaling, ferroptosis, ER protein processing, and apoptosis (**Supplementary Table S5**). The p53 signaling pathway showed significant enrichment in both treatments (ABT-751: 35 genes, p = 2.0×10⁻⁷; helenalin: 52 genes, p = 2.3×10⁻⁴), but with markedly different response magnitudes. Helenalin showed strong activation of DNA damage-related p53 activity and pro-apoptotic gene signaling, including *GADD45A*/*B*/*G* (3.1/5.4/4.7 log_2_-fold increase, respectively) [35], *FOS* (6.4 log_2_-fold increase) [36], and *DDIT3*/*CHOP* (3.3 log_2_-fold increase) [37]. This coincided with suppression of DNA repair and survival mechanisms such as *TP73* (1.6 log_2_-fold decrease) [38] and *PRKDC* (2.2 log_2_-fold decrease) [39]. The narrow response of ABT-751 was much more moderate and constrained to cell cycle-related p53 activity. While ABT-751 engaged a subset of the same genes as helenalin, such as the *GADD45* family, it did so at 3-to 5-fold lower levels. The treatment also showed modest repression of checkpoint regulators such as *CCNG2* (0.33 log_2_-fold decrease), consistent with its mechanism of action as a tubulin inhibitor.

Ferroptosis pathway enrichment was also significant in both treatments (ABT-751: 21 genes, p = 1.5×10⁻⁵; helenalin: 29 genes, p = 6.1×10⁻³), but the underlying gene expression patterns revealed two distinct stress phenotypes. *HMOX1* exhibited 3.8 log_2_-fold upregulation under helenalin versus 0.78 log_2_-fold under ABT-751, and *FTL* showed 3.5 log_2_-fold versus 0.71 log_2_-fold increases. Both genes are oxidative stress responders; a surge of both indicates a drive into ferroptotic damage. For helenalin, these markers were not accompanied by an adaptation to regenerate glutathione from cystine import (*SLC7A11* change n.s. [40]), thrusting the cell into a state with high ROS and low buffering capacity consistent with the agent’s thiol-binding mechanism. As with the p53 pathway, ABT-751 presented a much more moderate effect. While pro-ferroptotic expression of *HMOX1* and *FTL* increased mildly compared to DMSO, there was a small compensatory upregulation of *SLC7A11* (0.48 log-fold increase) with this treatment, indicating an adaptive response to maintain glutathione pools. Thus, while helenalin-induced changes overflowed into ferroptosis, ABT-751 promoted a protective response to ROS.

Most notably, helenalin induced significantly stronger ER protein processing pathway enrichment (132 genes, p = 1.3×10⁻¹⁴) compared to ABT-751 (59 genes, p = 7.2×10⁻⁶). A curated ER-stress/UPR gene panel (**Supplementary Table S6**) showed robust induction under helenalin of key markers including *DDIT3*/*CHOP*, *ATF4*, *HSPA5*/*BiP*, and *ERN1*/*IRE*1, whereas ABT-751 produced only modest increases in these markers. This indicates that helenalin triggered an extreme UPR response associated with cellular dysfunction; this could be mechanistically explained by rampant protein-misfolding as a result of helenalin’s reaction with cysteine. ABT-751, a mitotic rather than proteotoxic agent, maintained adaptive ER homeostasis.

Direct comparison of KEGG enrichment between treatments (**Fig. 6E**) showed that most pathways clustered along the diagonal, indicating similar overall activation levels. However, five pathways showed substantial divergence below the diagonal, demonstrating stronger enrichment in ABT-751-treated cells: Human T-cell leukemia virus 1 infection, viral carcinogenesis, p53 signaling pathway, apoptosis, and ferroptosis. Importantly, these pathways primarily capture host checkpoint and DNA-damage response mechanisms rather than pathogen-specific processes, reflecting ABT-751’s focused stress response program. These data indicate that although ABT-751 showed a higher ratio of enriched genes in these five areas, the magnitude of gene-level induction was lower, and these profiles show a concerted effort to counteract stress, rather than overflow into unproductive cell death.

The cell cycle pathway exhibited comparable enrichment with both agents, reinforcing that arrest alone cannot explain the productivity phenotype. However, expression distribution analysis (**Fig. 6F**) revealed that across all major pathways, helenalin consistently triggered broader and more extreme log₂ fold-changes compared to ABT-751’s controlled responses. This pattern indicates that while both treatments engage similar pathway networks, helenalin’s excessive transcriptional disruption likely overwhelms cellular adaptive capacity, whereas ABT-751 maintains a moderate stress activation sufficient to enhance biosynthetic processes without triggering uncontrolled cascades.

## 4 Discussion

### Divergent Effects of Cell-Cycle Inhibitors on rAAV Production

We investigated how two mechanistically distinct cell-cycle inhibitors, ABT-751, a microtubule-destabilizing agent, and helenalin, a thiol-reactive sesquiterpene lactone targeting NF-κB, impact recombinant AAV production in HEK293-derived systems [41], [42]. While both compounds successfully induced G₂/M arrest (ABT-751: 29% G₂/M; helenalin: 25% G₂/M; vs. 14% control; **Fig. 4B**), their effects on AAV yield diverged dramatically. ABT-751 consistently enhanced rAAV9 titers approximately 3-fold over DMSO controls and increased specific productivity by over 10-fold (**Fig. 4C**), while helenalin severely suppressed vector output below control levels. These findings demonstrate that cell-cycle arrest alone is insufficient to predict rAAV productivity; rather, the specific cellular programs engaged by each compound determine the outcome.

### Cell-Cycle Position Cannot Explain Productivity Differences

Analysis of specific productivity revealed that ABT-751 enhances per-cell viral output by approximately 5-fold compared to dividing controls. This confirms that the titer increase is driven by a fundamental upregulation of cellular manufacturing capacity, independent of the biomass reduction associated with G₂/M arrest (**Fig. 4D**).

### Adaptive versus Maladaptive Stress Responses Determine Productivity

Our transcriptomic analysis reveals fundamental differences in how G_2_/M-arresting compounds engage cellular stress pathways. Principal component analysis showed that helenalin-treated samples separated dramatically from both control and ABT-751 groups along PC1 (81% of variance, **Fig. 5A**), with GSEA revealing this axis primarily reflects suppression of mitochondrial energy metabolism and activation of severe stress responses (**Supplementary Table S1**).

The scale of transcriptional disruption differed markedly: helenalin dysregulated 4,579 annotated genes compared to 1,895 for ABT-751. Hierarchical clustering of the 50 most variable genes revealed distinct expression modules specific to each treatment. ABT-751 induced what we characterize as productive stress; moderate activation of checkpoint genes without triggering severe apoptotic programs. Notably, ABT-751 did not significantly elevate *DDIT3/CHOP* expression, a key pro-apoptotic UPR effector [43], [44], and maintained cell viability above 65% throughout the production period.

In contrast, helenalin triggered excessive stress responses characterized by dramatically stronger gene activation. Key stress markers in p53, apoptotic, and ferroptotic pathways showed 2-5 log_2_-fold greater induction under helenalin – for example, *GADD45A*: 3.1 log_2_-fold vs. 0.52 log_2_-fold for ABT-751; *HMOX1*: 3.8 log_2_-fold vs. 0.78 log_2_-fold, and *SERPINE1*: 5.5 log_2_-fold vs. 2.2 log_2_-fold (**Supplementary Table S6**). Together, this indicates activation of cell death pathways and shutdown of repair machinery. The robust induction of *CHOP* and comprehensive UPR activation under helenalin indicates a shift from adaptive to terminal, maladaptive stress responses that compromise biosynthetic function [45]. Generally, across stress pathways, helenalin demonstrated a runaway effect characterized by immense swings in gene expression, yielding an irrecoverable state with lower cell productivity. Alternatively, ABT-751 also affected a significant number of pathways, but with less severe individual gene expression and compensatory responses that worked to restore homeostasis, such as through relief of oxidative stress.

### Focused versus Dispersed Transcriptional Programs

KEGG pathway enrichment comparison revealed a key distinction in the focus of the transcriptional responses between the two treatments. Helenalin induced a broad, high-magnitude transcriptional response, as visualized by the wide gene expression distributions (**Fig. 6F**) and the large number of total mapped DEGs (4,579). Because this impact was so dispersed, helenalin’s top enriched KEGG pathways were dominated by broad cellular processes like “Endocytosis” and “Protein processing in endoplasmic reticulum” (**Fig. 6D**). Notably, although genes within key stress pathways (e.g., p53, Apoptosis, Ferroptosis) were strongly impacted by helenalin at the gene level (**Fig. 6F**), these pathways were not among the top-ranked statistical enrichments, likely masked by the massive, system-wide disruption consistent with an integrated stress response [46]. In sharp contrast, ABT-751 induced a more confined transcriptional response, characterized by moderate fold-changes (**Fig. 6F**) and far fewer DEGs (1,895). Due to this focused impact, these same pathways—”p53 signaling,” “Apoptosis,” and “Ferroptosis”—were highly ranked as statistically significant enrichments for ABT-751 (**Fig. 6C**). This comparison demonstrates that ABT-751 elicited a focused, mild response on these key pathways, whereas helenalin triggered a stronger, more dispersed transcriptional cascade.

The enrichment of neurodegeneration pathways uniquely in helenalin-treated cells (amyotrophic lateral sclerosis, spinocerebellar ataxia) further suggests proteotoxic stress and cellular dysfunction incompatible with productive rAAV assembly. Expression distribution analysis across pathways confirmed that helenalin consistently triggered broader and more extreme log₂ fold-changes compared to ABT-751’s focused responses. Meanwhile, both treatments showed comparable cell cycle pathway enrichment, reinforcing that cell cycle arrest magnitude alone cannot explain productivity differences.

### Platform Applicability, Manufacturing Implications, and Agent Selection

Importantly, ABT-751’s benefits extended across multiple AAV serotypes (AAV2: 4.7-fold increase; AAV5: 6.6-fold; AAV9: 4.5-fold) and production platforms, with even greater enhancements observed in Cell Line A compared to Expi293F. These consistent improvements across diverse systems suggest ABT-751 engages fundamental cellular mechanisms favorable for AAV production. Furthermore, our selected protocol involved low concentrations of ABT-751 (0.97 µM) following transfection, balancing cell-cycle modulation with cost and time savings.

The Venn diagram analysis (**Fig. 5F**) revealed 2,833 genes commonly dysregulated across all contrasts (core stress response), but also distinct gene sets unique to each compound (237 for ABT-751, 2,065 for helenalin), suggesting that compounds can be rationally selected based on their stress response profiles rather than simple arrest efficiency. However, the proper approach can be further elucidated through additional dynamics of stress response activation over extended periods to determine whether helenalin’s maladaptive responses could be limited through a more optimized timing of chemical addition. Furthermore, while we demonstrated titer improvements, additional quality attributes including empty/full capsid ratios, genome integrity, and in vivo potency should be assessed to further establish the value of these chemical inducer compounds in a biomanufacturing process.

### Mechanistic Insights and Cell Engineering Opportunities

Our data support a model where productive AAV manufacturing requires not just cell-cycle synchronization but also maintenance of cellular homeostasis during arrest. The initial screening of multiple cell-cycle inhibitors revealed that compounds achieving similar G₂/M enrichment (ABT-751, helenalin) produced vastly different titer outcomes, emphasizing the importance of compound-specific cellular responses.

Further, the quantitative expression patterns identified here suggest rational targets for cell line engineering. Genes showing controlled activation under productive ABT-751 conditions (e.g., moderate enhancement of iron homeostasis genes like *FTL* with concurrent enhancement of *SLC7A11*), ER chaperones like *HSPA5*/*BiP*) could be targeted for stable expression, while pro-apoptotic effectors like CHOP could be knocked down to prevent uncontrolled responses. AAV Rep proteins interact directly with cell cycle regulators including p53 [48] and pRb [49], providing a mechanistic link between cell cycle modulation and AAV production. These interactions may explain why G₂/M arrest specifically benefits AAV production, as Rep proteins can leverage the arrested state to enhance viral genome replication and packaging.

Although subsets of pathways were more enriched in ABT-751-treated cultures, gene-level analyses of potential engineering targets suggested lower individual gene induction and a transcriptional rebalancing to moderate stresses. This aligns with the concept that mild stress adjustments can enhance cellular resilience and productivity [47].

## 5 Conclusion

Our findings demonstrate that improving rAAV production through cell-cycle intervention depends critically on maintaining cellular homeostasis during arrest. ABT-751 achieves productive G₂/M blockade while preserving biosynthetic capacity. Helenalin, despite similar G_2_/M arrest, provokes dysregulated stress responses that ultimately undermine vector production. These findings establish a framework for rational selection of cell cycle modulators based on their stress response profiles rather than simple arrest efficiency and highlight potential future cell engineering targets. Together, these process improvements expand and enhance the capabilities of current and next-generation AAV manufacturing platforms, working towards reduced costs for gene therapies of the future.

## Data Availability Statement

The data that support the findings of this study are available from the corresponding author upon reasonable request.

## Acknowledgements

The authors would like to thank Hanhvy Bui (Integrated Imaging Center, Johns Hopkins University) for assistance with flow cytometry studies.

## Author Contributions

**J.W.:** Conceptualization, data curation, formal analysis, investigation, methodology, software, validation, writing – original draft, writing – review and editing; **J.S.:** Conceptualization, formal analysis, investigation, methodology, software, validation, writing – original draft, writing – review and editing; **D.C.:** Formal analysis, investigation, validation, writing – review and editing; **N.G.:** Formal analysis, investigation, validation, writing – original draft, writing – review and editing; **M.J.B.:** Conceptualization, funding acquisition, methodology, project administration, supervision, validation, writing – original draft, writing – review and editing.

