## Supplemental Figures and Captions for "Tailored Cell Cycle Modulation Enhances AAV Manufacturing: Balancing Arrest with Adaptive Stress Responses"

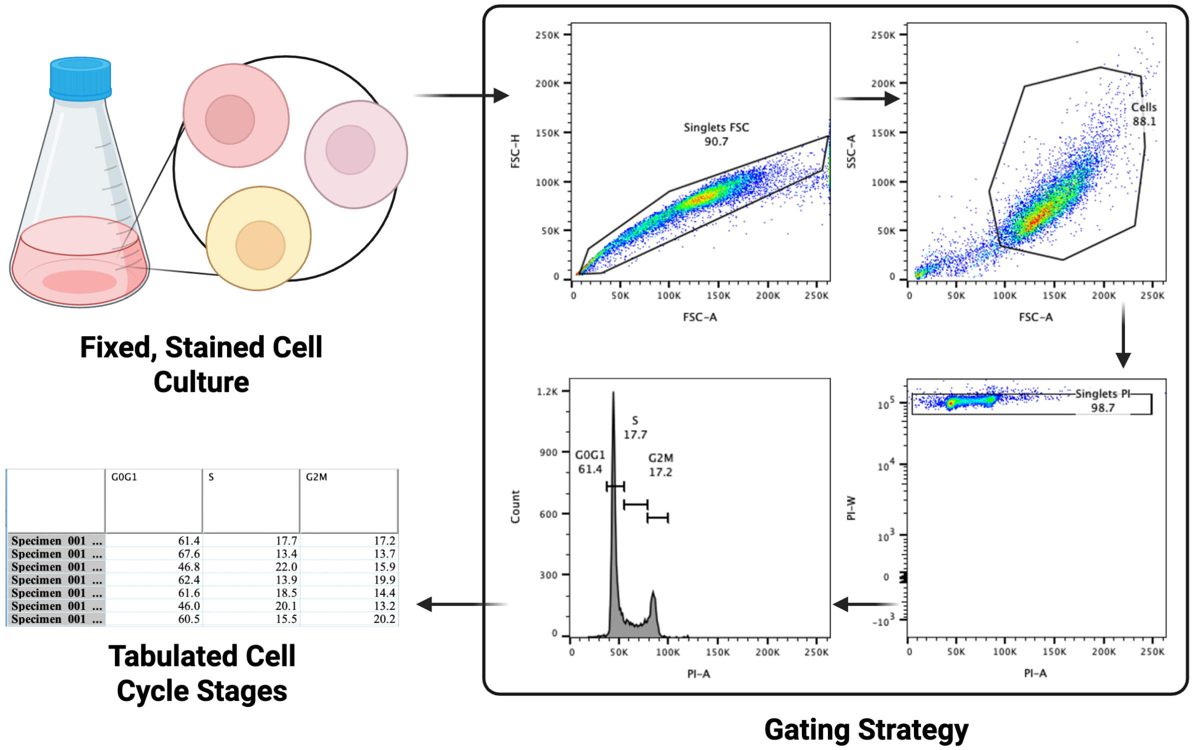


**Figure S1: Flow cytometry workflow and gating for PI-based cell-cycle profiling.** Schematic (left) shows sampling of cells from culture for DNA-content analysis. Representative flow cytometry plots (right) illustrate the sequential gating workflow: (i) singlet discrimination using forward scatter (FSC), (ii) cell gate on FSC/SSC to exclude debris, and (iii) PI-positive singlets used for DNA-content measurement. The cell-cycle stage assignments were made by visual inspection of the PI histogram to estimate $G_1$, S, and $G_2$/M fractions. The table summarizes phase distributions across replicate samples. Percentages displayed on the plots are from a representative example. Abbreviations: FSC, forward scatter; SSC, side scatter; PI, propidium iodide. Created with BioRender.

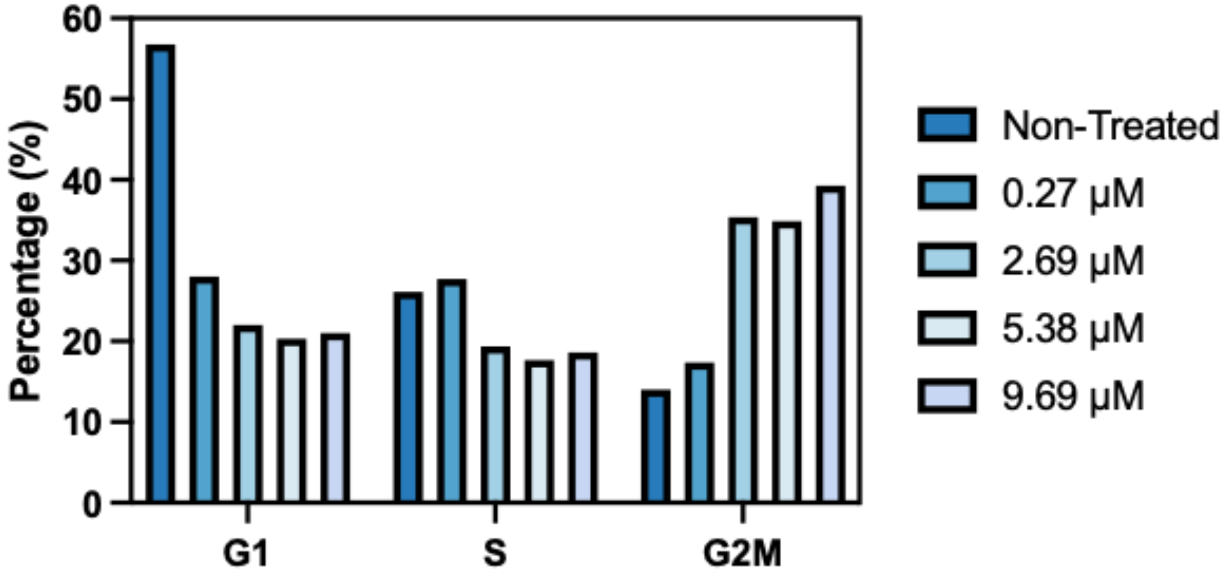


**Figure S2: Effect of various ABT-751 concentrations on AAV2 cell cycle distribution.** Quantification of cell cycle distribution showing the percentage of cells in different phases (G1, S, and G2/M) three days post-transfection. HEK293 cells were either non-treated or treated with ABT-751 at 0.269 μM, 2.69 μM, 5.38 μM, or 9.69 μM. Data are presented as value of *n* = 1 replicate.

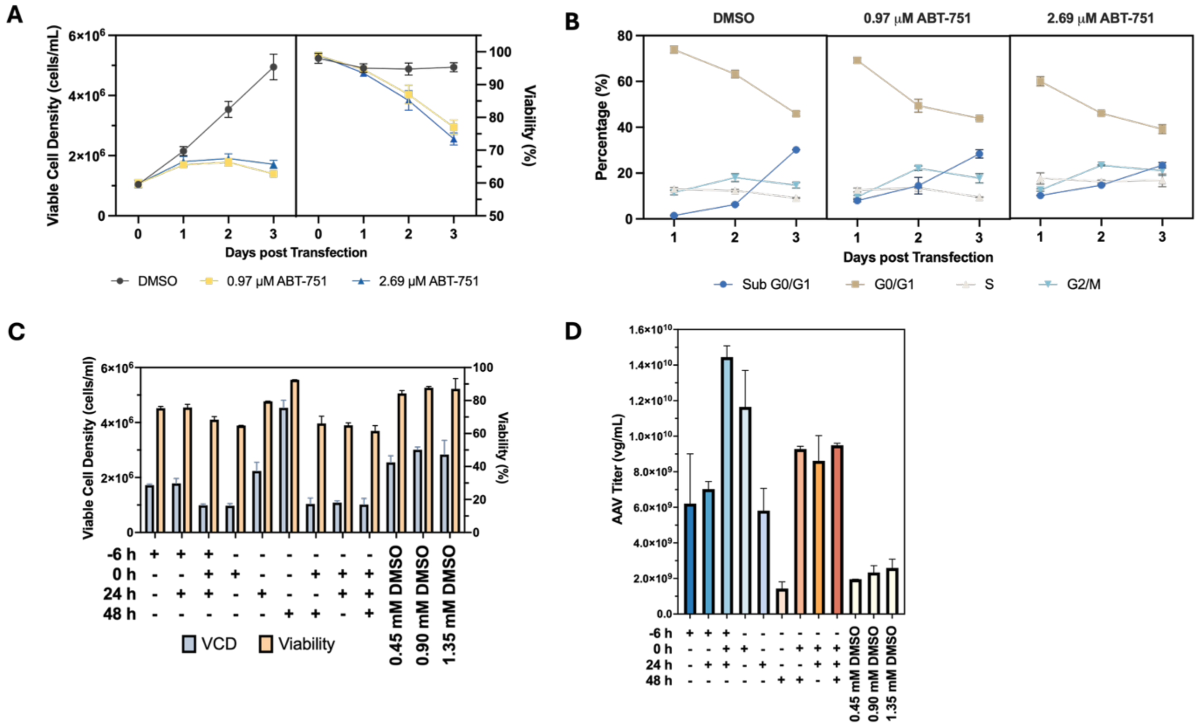


**Figure S3: Effect of ABT-751 treatment and spiking strategies on AAV2 production and cell viability. (A)** Viable cell density and viability for HEK293 cells producing AAV treated with DMSO, 0.97 µM ABT-751, or 2.69 µM ABT-751. Data are presented as mean ± SD (*n* = 3 independent biological replicates). **(B)** Quantification of cell cycle distribution showing the percentage of cells in different phases (Sub-G$_1$, G$_1$, S, and G$_2$/M) over three days post-transfection. HEK293 cells were treated with either vehicle (DMSO) or ABT-751 at two concentrations (2.69 µM and 0.97 µM). Data are presented as mean ± SD (*n* = 3 independent biological replicates). **(C)** Viable cell density (VCD, million/mL) and viability (%) across different spiking strategies. ABT-751 (0.97 µM) was added at the indicated time points: 6 hours before transfection (-6 h), at transfection (0 h), 24 hours after transfection (24 h), and 48 hours after transfection (48 h). DMSO was added at 0 h of transfection. Individual values are plotted with SD (*n* = 2 independent biological replicates). **(D)** AAV2 titer (vector genomes per mL, vg/mL) across different spiking strategies. Labels indicate spiking conditions, with different time points and DMSO concentrations. Data are presented as mean ± SD (*n* = 2 independent biological replicates).

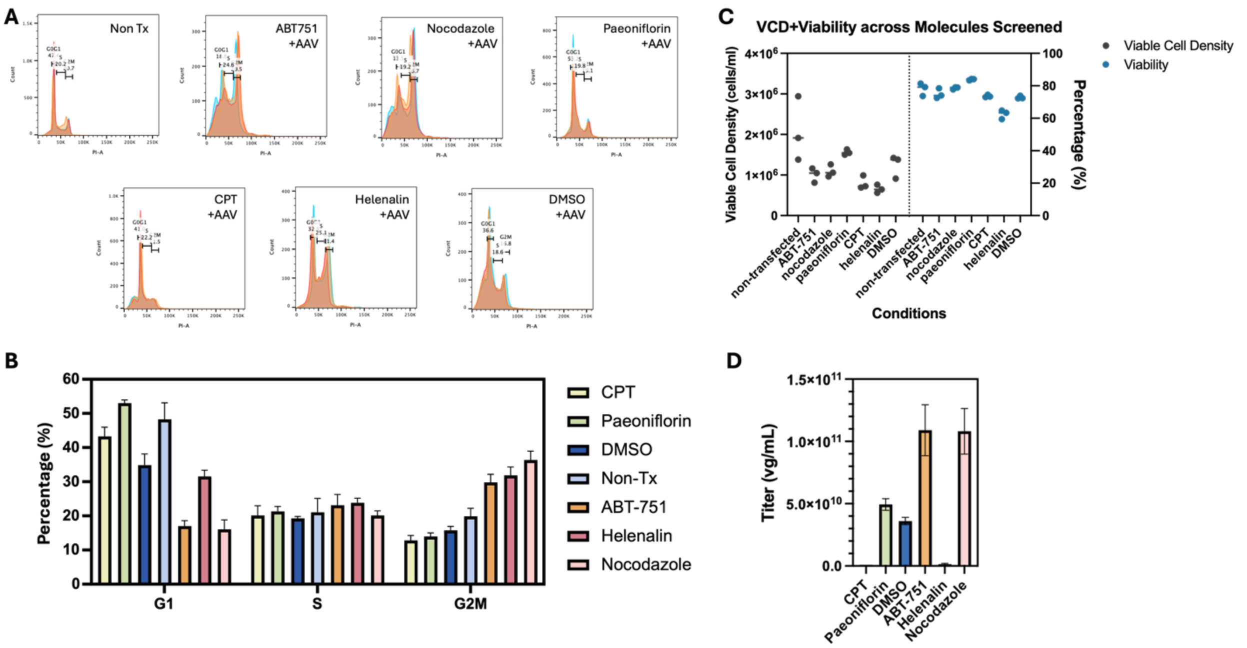


**Figure S4: Cell-cycle and rAAV titer effects of candidate molecules in Expi293 cells.** **(A)** Representative day-3 PI DNA-content histograms for the indicated conditions (Non-Tx, DMSO, 0.97 µM ABT-751, 4 µM nocodazole, 500 µM paeoniflorin, 20 µM CPT, 5 µM helenalin). **(B)** Summary of cell-cycle distributions across replicates shown as stacked bars ($G_1$, S, $G_2$/M). Bars show mean ± SD (*n=3*). **(C)** Viable cell density (VCD) and viability (%). Each point is one biological replicate (well). **(D)** Bulk rAAV vector-genome titer (vg/mL) at day 3, mean ± SD across wells (*n=3*).

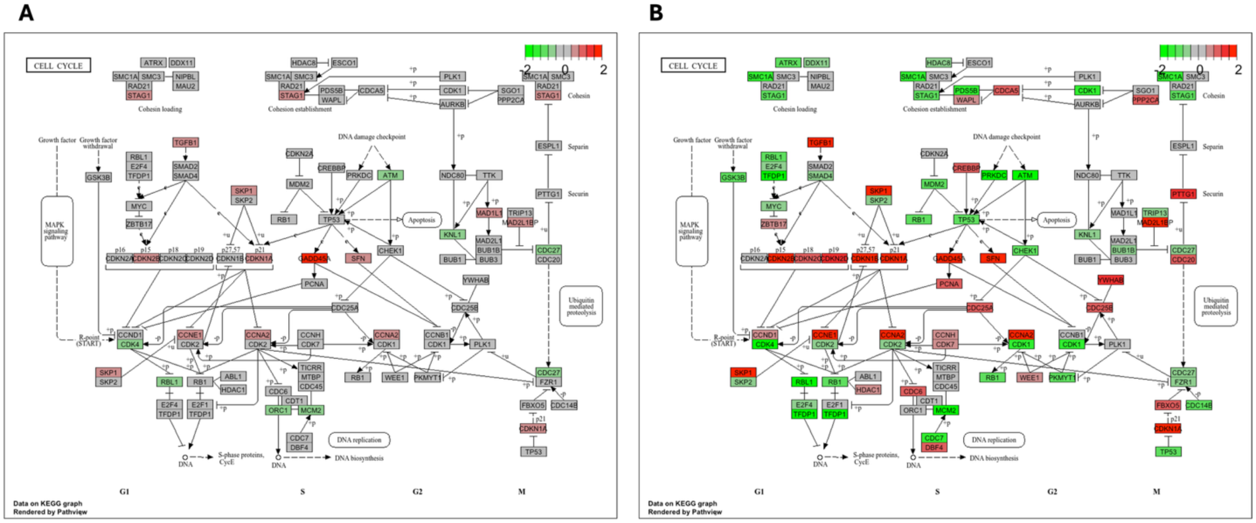


ABT-751 vs DMSO　　　　Helenalin vs DMSO

**Supplementary Figure S5. KEGG Cell Cycle Pathway Analysis Reveals Differential Regulatory Responses to ABT-751 and Helenalin Treatment. (A)** Cell cycle pathway map showing differential gene expression in ABT-751-treated versus DMSO control HEK293 cells at 72 hours post-transfection. **(B)** Cell cycle pathway map showing differential gene expression in helenalin-treated versus DMSO control cells under identical conditions. Gene expression changes are visualized using a color gradient where red indicates upregulation, green indicates downregulation, and gray indicates no significant change (padj ≥ 0.05). Color intensity corresponds to the magnitude of $\log_2$ fold-change. Both compounds induce $G_2$/M arrest but with distinct regulatory patterns: helenalin shows stronger activation of G2/M checkpoint kinases (WEE1, CHEK1) and more complete suppression of the APC/C complex (CDC20, CDC27) required for mitotic exit, while ABT-751 displays more selective effects with retention of partial S/G2 cyclin expression (CCNA2) and $G_1$/S cyclins (CCNE1). The pathway visualization was generated using Pathview with differential expression data from RNA-seq analysis. Key cell cycle phases are indicated ($G_1$, S, $G_2$, M), along with major regulatory complexes including the MAPK signaling pathway input, growth factor withdrawal signals, DNA damage checkpoints, and ubiquitin-mediated proteolysis. These distinct expression patterns demonstrate that while both compounds arrest cells at $G_2$/M, helenalin enforces multiple checkpoint barriers more stringently than ABT-751, consistent with its broader transcriptional disruption.
